# Incorporation of doxorubicin in different polymer nanoparticles and their anti-cancer activity

**DOI:** 10.1101/403923

**Authors:** S. Pieper, H. Onafuye, D. Mulac, Jindrich Cinatl, Mark N. Wass, M. Michaelis, K. Langer

## Abstract

Nanoparticles are under investigation as carrier systems for anti-cancer drugs. They have been shown to accumulate in cancer tissues through the enhanced permeability and retention (EPR) effect, to reduce toxicity to non-target tissues, and to protect drugs from preliminary inactivation. However, nanoparticle preparations are not commonly compared for their anti-cancer effects at the cellular level. Here, we prepared doxorubicin-loaded nanoparticles based on poly(lactic-*co*-glycolic acid) (PLGA), polylactic acid (PLA), and PEGylated PLGA (PLGA-PEG) by solvent displacement and emulsion diffusion approaches. The resulting nanoparticles covered a size range between 73 and 246 nm. PLGA-PEG nanoparticle preparation by solvent displacement resulted in the smallest nanoparticles. In PLGA nanoparticles, the drug load could be optimised using solvent displacement at pH7 reaching 53 µg doxorubicin/mg nanoparticle. In addition, these PLGA nanoparticles displayed sustained doxorubicin release kinetics compared to the more burst-like kinetics of the other preparations. In neuroblastoma cells, doxorubicin-loaded PLGA-PEG nanoparticles (presumably due to their small size) and PLGA nanoparticles prepared by solvent displacement at pH7 (presumably due to their high drug load and superior drug release kinetics) exerted the strongest anti-cancer effects. In conclusion, doxorubicin-loaded nanoparticles made by different methods from different materials displayed substantial discrepancies in their anti-cancer activity at the cellular level. Optimised preparation methods resulted in PLGA nanoparticles characterised by increased drug load, controlled drug release, and high anti-cancer efficacy. The design of drug-loaded nanoparticles with optimised anti-cancer activity at the cellular level is an important step in the development of improved nanoparticle preparations for anti-cancer therapy.

## INTRODUCTION

According to Globocan, there “were 14.1 million new cancer cases, 8.2 million cancer deaths and 32.6 million people living with cancer (within 5 years of diagnosis) in 2012 worldwide” [http://globocan.iarc.fr/Pages/fact_sheets_cancer.aspx]. Despite substantial improvements over recent decades, the prognosis for many cancer patients remains unacceptably poor. In particular, the outlook is grim for patients that are diagnosed with disseminated (metastatic) disease who cannot be successfully treated by local treatment (surgery, radiotherapy). These patients depend on systemic drug therapy. However, the therapeutic window is small, and anti-cancer therapies are typically associated with severe side-effects [Steeg, 2016; Siegel et al., 2018].

One strategy to develop more effective cancer therapies is to use nano-sized drug delivery systems that mediate a more specific tumour accumulation of transported drugs. Tumour targeting can be achieved via the enhanced permeability and retention (EPR) effect, which is the consequence of increased leakiness of the tumour vasculature and a lack of lymph drainage [Rodallec et al., 2018]. Nano-sized drug carrier systems can also prolong the circulation time of anti-cancer drugs, protect them from degradation, and sustain therapeutic drug concentrations due to prolonged/ controlled drug release. In addition, nanoparticles can be used to administer poorly soluble agents, as demonstrated for Nab-paclitaxel, an HSA nanoparticle-based paclitaxel preparation approved for the treatment of different forms of cancer [Brufsky, 2017; Mir et al., 2017; Ricciardi et al., 2018; Rodallec et al., 2018; Tan et al., 2018; Zhao et al., 2018].

Another important aspect of the efficacy of nanoparticles as delivery system for anti-cancer is their uptake and, in turn, the drug transport into cancer cells. Uptake mechanisms may differ between different types of nanoparticles, which may affect their effectiveness as carriers for anti-cancer drugs. Here, we prepared and directly compared the effects of doxorubicin-loaded polylactic acid (PLA) and poly(lactic-*co*-glycolic acid) (PLGA) nanoparticles in neuroblastoma cells. PLA and PLGA are FDA- and EMA-approved for human use [Wischke & Schwendeman, 2008; Tyler et al., 2016] and are easily degraded into their monomers, lactic acid and glycolic acid. Furthermore, a copolymer composed of polyethylene glycol (PEG) and PLGA (PLGA-PEG) was used for nanoparticle preparation. PEGylated (“stealth”) nanoparticles display prolonged systemic circulation time, because they avoid agglomeration, opsonisation, and phagocytosis [Suk et al., 2016]. Doxorubicin was incorporated into nanoparticles prepared from these three polymers by emulsion diffusion or solvent displacement approaches. The resulting nanoparticles were compared by particle diameter, polydispersity index, zeta potential, drug load, and drug release behaviour. Selected preparations were tested for anti-cancer efficacy in cell culture.

## MATERIALS AND METHODS

### Reagents and chemicals

PLGA (Resomer^®^RG502H), PLA (Resomer^®^R203H) and PLGA-PEG (Resomer^®^ RGP d 50155) were obtained from Evonik Industries AG (Essen, Germany). Ethyl acetate, dichloromethane and methanol were purchased from VWR International GmbH (Darmstadt, Germany). Acetone, acetonitrile and dimethyl sulfoxide (DMSO) were obtained from Carl Roth GmbH (Karlsruhe, Germany). Poly(vinyl alcohol) (PVA, 30,000–70,000 Da), bovine serum albumin (BSA), HSA, and glutaraldehyde were obtained from Sigma-Aldrich Chemie GmbH (Karlsruhe, Germany). Dulbecco’s Phosphate buffered saline (PBS) was purchased from Biochrom GmbH (Berlin, Germany). Doxorubicin was obtained from LGC Standards GmbH (Wesel, Germany). All chemicals were of analytical grade and used as received.

### Nanoparticle preparation via emulsion diffusion

PLA and PLGA nanoparticles were prepared by a previously described emulsion diffusion technique [Michaelis et al., 2000; Astete & Sabliov, 2006]. PLA, PLGA, PLGA-PEG were dissolved in organic solvents (Table 1) and 200 µL of a methanolic doxorubicin solution (2.5 mg/mL) was added. This solution was then poured into 5 mL (1%, m/v) PVA solution and afterwards homogenized with an Ultra Turrax (IKA-Werke, Staufen, Germany) as indicated in Table 1. Subsequently this pre-emulsion was mixed with another 5 mL (1%, m/v) PVA solution. After stirring overnight, the resultant nanoparticles were purified three times by centrifugation at 21,000 g for 15 min (Eppendorf Centrifuge 5430 R, Eppendorf, Hamburg, Germany) and re-dispersion in purified water.

**Table 1:** Preparation parameters for nanoparticles based on different polymers using emulsion diffusion technique.

| Polymer | Amount of polymer | Organic solvent | Homogenisation |
| --- | --- | --- | --- |
| PLGA | 50 mg | 2.5 mL ethyl acetate | 15,000 g for 5 min |
| PLA | 100 mg | 2.0 mL dichloromethane | 18,000 g for 15 min |
| PLGA-PEG | 50 mg | 2.5 mL ethyl acetate | 15,000 g for 5 min |

After the final purification step, an aliquot of the nanoparticle suspension was centrifuged and the resulting pellet was dissolved in 1 mL DMSO in order to measure the entrapped amount of doxorubicin by HPLC (see below).

In order to increase the drug load for PLGA nanoparticles different volumes of the methanolic doxorubicin solution (2.5 mg/mL) were used corresponding to 1.0, 1.5, and 2.0 mg total doxorubicin. For a further increase in drug load different aqueous doxorubicin solutions (ranging from 10.0 to 50.0 mg/mL) were used to achieve total doxorubicin amounts of 0.5, 2.5, 5.0, 7.5, 12.5, and 25.0 mg. In both cases the amount of the polymer was kept constant at 50 mg. To prepare doxorubicin-loaded nanoparticles at a defined pH of 7, a PVA solution (1%, m/v) in phosphate buffer (15.6 mg/mL NaH_2_PO_4_ • 2 H_2_O; pH adjusted to 7 with NaOH) was used.

### Nanoparticle preparation via solvent displacement

For nanoparticle preparation via solvent displacement technique 60 mg polymer were dissolved in 2 mL acetone and combined with 200 µL doxorubicin solution (2.5 mg/mL). This mixture was injected into 4 mL 2% (m/v) PVA solution to produce PLGA and PLGA-PEG nanoparticlesor into 4 mL 1% (m/v) PVA solution to produce PLA nanoparticles. After stirring overnight at 550 rpm and evaporation of the organic solvent, PLA and PLGA nanoparticles were purified three times by centrifugation at 21,000 g for 15 min and re-dispersion in purified water. PLGA-PEG nanoparticles were purified three times by centrifugation at 30,000 g for 60 min and re-dispersion in purified water.

### Determination of particle size, size distribution and zeta potential

Average particle size and the polydispersity were measured by photon correlation spectroscopy (PCS) using a Malvern zetasizer nano (Malvern Instruments, Herrenberg, Germany). The resulting particle suspensions were diluted 1:100 with purified water and measured at a temperature of 22°C using a backscattering angel of 173°. The zeta potential was determined with the same instrument and the same diluted nanoparticle suspension by Laser Doppler microelectrophoresis.

### Scanning electron microscopy (SEM)

For scanning electron microscopy (SEM), the particle suspensions were diluted with purified water to 0.25 mg/mL. The suspension was dropped on a filter (MF-Millipore™ membrane filter VSWP, 0.1 µm) and dried for 24 h in a desiccator. Afterwards, the membranes were sputtered with gold under argon atmosphere (SCD 040, BAL-TEC, Balzers, Liechtenstein). The SEM pictures were received at an accelerating voltage of 10,000 V and a working distance of 10 mm (CamScan CS4, Cambridge Scanning Company, Cambridge, UK).

### Doxorubicin quantification via HPLC-UV

The amount of doxorubicin that had been incorporated incorporated into the nanoparticles was determined by HPLC-UV (HPLC 1200 series, Agilent Technologies GmbH, Böblingen, Germany) using a LiChroCART 250 x 4 mm LiChrospher 100 RP 18 column (Merck KGaA, Darmstadt, Germany). The mobile phase was a mixture of water and acetonitrile (70:30) containing 0.1% trifluoroacetic acid [Dreis et al., 2007]. In order to obtain symmetric peaks a gradient was used. In the first 6 min the percentage of water was reduced from 70% to 50%. Subsequently within 2 min the amount of water was further decreased to 20% and then within another 2 min increased again to 70%. These conditions were hold for a final 5 min resulting in a total runtime of 15 min. While using a flow rate of 0.8 mL/min, an elution time for doxorubicin of t = 7.5 min was achieved. The detection of doxorubicin was performed at a wavelength of 485 nm [Sanson et al., 2010].

### *In vitro* drug release studies

To study drug release *in vitro*, a nanoparticle suspension of 1 mg nanoparticles in 1 mL of PBS containing 5% (m/v) bovine serum albumin (BSA) was shaken at 37°C with 500 rpm. Nanoparticle suspensions were centrifuged (30,000 g, 15 min) after 0, 0.5, 1, 2, 4, 6, 8, and 24 h, and an aliquot (250 µL) of the supernatant was diluted with 750 µL ethanol (96%, v/v) in order to precipitate BSA. After a second centrifugation step (30,000 g, 10 min) the supernatant was analysed for the amount of released doxorubicin as mentioned above. Additionally, the resulting pellet was dissolved in DMSO in order to calculate doxorubicin recovery.

### Cell culture

The MYCN-amplified neuroblastoma cell line UKF-NB-3 was established from stage 4 neuroblastoma patients [Kotchetkov et al., 2005]. UKF-NB-3 sub-lines adapted to growth in the presence of doxorubicin 20ng/mL (UKF-NB-3^r^DOX^20^) [Kotchetkov et al., 2005] or vincristine 1ng/mL (UKF-NB-3^r^VCR^1^) were established by continuous exposure to step-wise increasing drug concentrations as described previously described [Kotchetkov et al., 2005; Michaelis et al., 2011] and derived from the Resistant Cancer Cell Line (RCCL) collection (https://research.kent.ac.uk/ibc/the-resistant-cancer-cell-line-rccl-collection/).

All cells were propagated in Iscove’s modified Dulbecco’s medium (IMDM) supplemented with 10 % foetal calf serum, 100 IU/ml penicillin and 100 mg/ml streptomycin at 37°C. The drug-adapted sub-lines were continuously cultured in the presence of the indicated drug concentrations. Cells were routinely tested for mycoplasma contamination and authenticated by short tandem repeat profiling.

### Cell viability assay

Cellviabilitywasdeterminedby3-(4,5-dimethylthiazol-2-yl)-2,5-diphenyltetrazolium bromide (MTT) assay modified after Mosman [Mosman, 1983], as previously described [Michaelis et al., 2000]. 2×10^4^ cells suspended in 100 µL cell culture medium were plated per well in 96-well plates and incubated in the presence of various concentrations of drug or drug preparations for 120 h. Then, 25µL of MTT solution (1 mg/mL (w/v) in PBS) were added per well, and the plates were incubated at 37°C for an additional 4h. After this, cells were lysed using 200µL of a buffer containing 20% (w/v) sodium dodecylsulfate and 50% (v/v) N, N-dimethylformamide (pH 4.7) at 37°C for 4h. Absorbance was determined at 570 nm for each well using a 96-well multiscanner. After subtracting of the background absorption, the results are expressed as percentage viability relative to untreated control cultures. Drug concentrations that inhibited cell viability by 50% (IC50) were determined using CalcuSyn (Biosoft, Cambride, UK).

## RESULTS

### Influence of the preparation technique on particle diameter and polydispersity index

Particle diameters are presented in Figure 1. Emulsion diffusion (173.5 ± 5.9 nm) and solvent displacement (179.4 ± 7.6 nm) resulted in PLGA nanoparticles with similar diameters. In contrast, solvent displacement resulted in PLGA-PEG nanoparticles 72.6 ± 3.3 nm whereas emulsion diffusion resulted in PLGA-PEG nanoparticles > 200 nm. In accordance, solvent displacement using the stabiliser PVA at concentrations between 2-4% (w/v) and controlled injection at mild stirring had previously been shown to produce PLGA-PEG nanoparticles with a diameter < 100 nm [Kwon et al., 2001; Astete and Sabliov, 2006; Zhou et al., 2015]. The hydrophilic PEG chains may sterically stabilise the nanoparticles by reducing PLGA aggregation during nanoparticle formation resulting in smaller particle diameters [Ameller et al., 2003].

**Figure 1:**
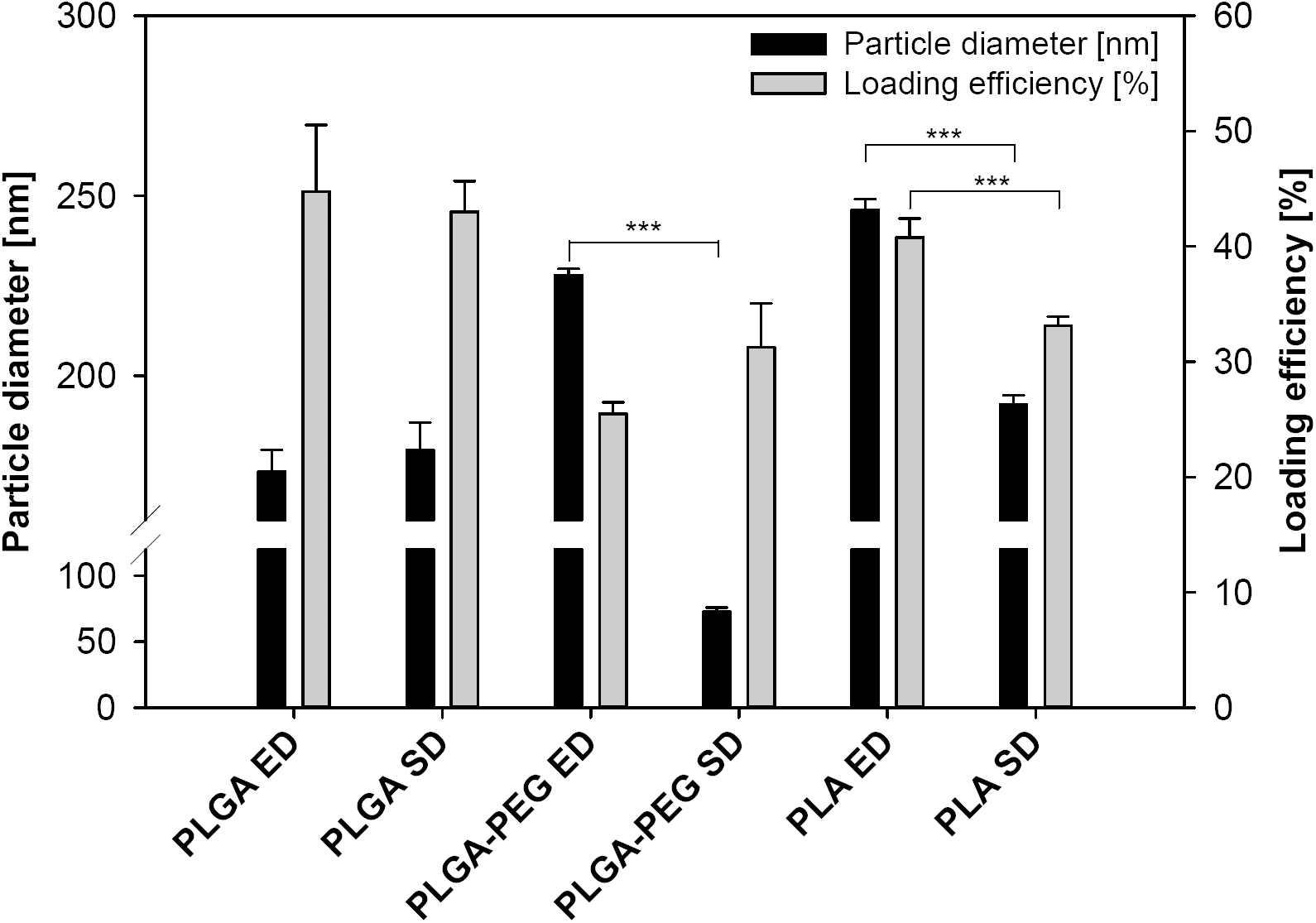
Resulting particle diameters and loading efficiencies for different nanoparticle mformulations using emulsion diffusion (ED) or solvent displacement (SD) techniques (data expressed as means ± SD, n ≥ 3).

Emulsion diffusion resulted in PLA nanoparticles of 246.2 ± 2.9 nm and solvent displacement in PLA nanoparticles of 192.1 ± 2.5 nm. Polydispersity indices < 0.1 indicated a monodisperse size distribution for all nanoparticle preparations. Monodispersity and particle diameters were confirmed by scanning electron microscopy (SEM) images (Figure 2).

**Figure 2:**
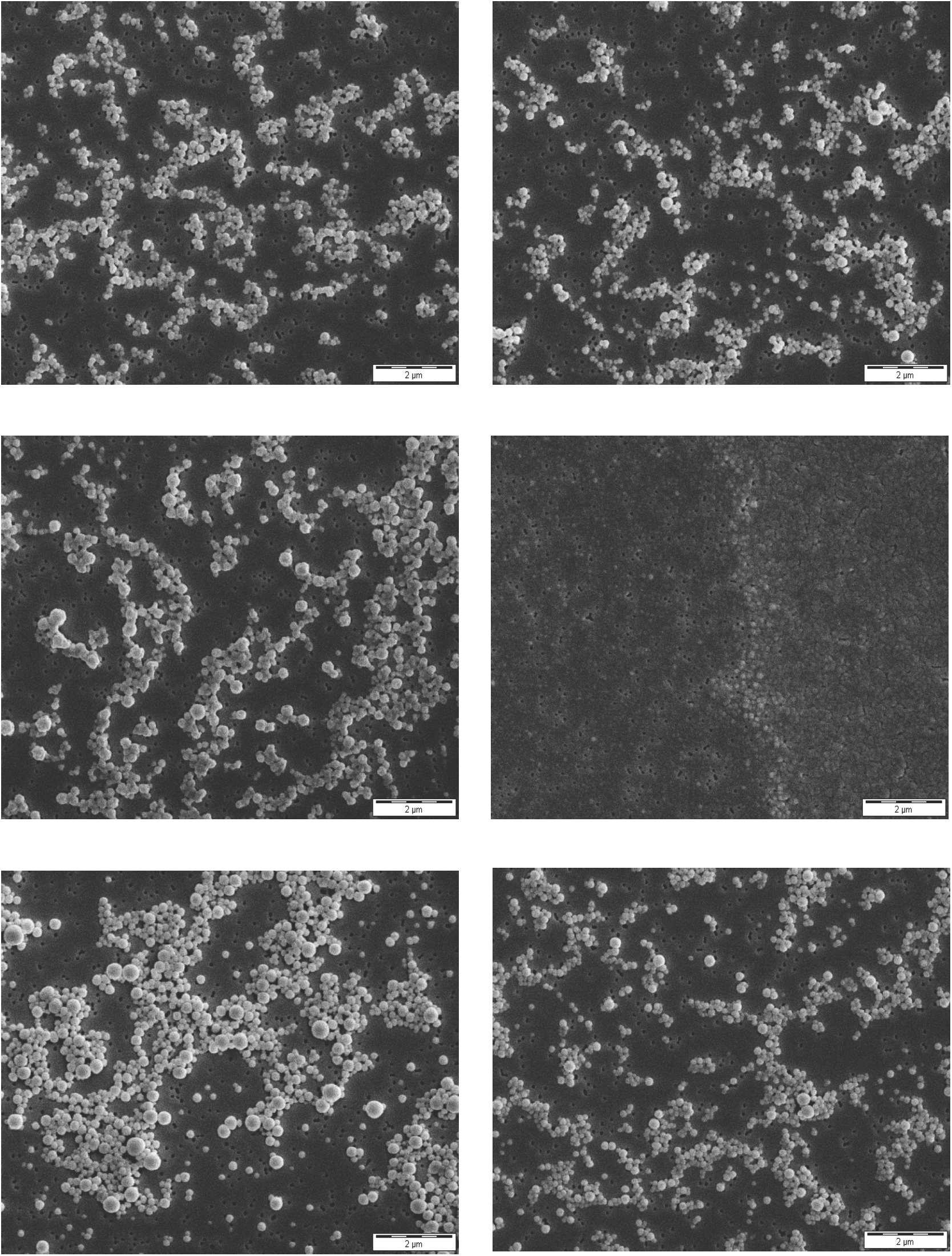
SEM images of nanoparticles using emulsion diffusion (ED) or solvent displacement (SD) preparation technique. (A) PLGA nanoparticles ED, (B) PLGA nanoparticles SD, (C) PLGA-PEG nanoparticles ED, (D) PLGA-PEG nanoparticles SD, (E) PLA nanoparticles ED, (F) PLA nanoparticles SD. Images were taken at 10,000x magnification.

### Influence of the preparation technique on loading efficiency and drug release

Loading efficiencies ranging from 25.5 ± 1.0% to 44.8 ± 5.8% of the applied doxorubicin were detected in the different nanoparticle preparations (Figure 1), resulting in drug loads between 2.6 ± 0.2 µg doxorubicin/mg nanoparticle and 6.7 ± 0.3 µg doxorubicin/mg nanoparticle (Table 2).

**Table 2:** Nanoparticle (NP) yield and doxorubicin (Dox) drug load results for nanoparticles prepared by either emulsion diffusion (ED) or solvent displacement (SD) technique (data expressed as means ± SD, n ≥ 3).

| NP system | NP yield | NP yield | Drug load |
| --- | --- | --- | --- |
| | [mg NP/mL] | [%] | [μg Dox/mg NP] |
| PLGA ED | $3.3 \pm 0.4$ | $66.8 \pm 7.2$ | $6.7 \pm 0.3$ |
| PLGA SD | $8.5 \pm 0.4$ | $70.4 \pm 3.0$ | $5.1 \pm 0.2$ |
| PLGA-PEG ED | $4.2 \pm 0.1$ | $84.4 \pm 1.8$ | $3.0 \pm 0.2$ |
| PLGA-PEG SD | $7.6 \pm 0.9$ | $63.6 \pm 7.4$ | $4.1 \pm 0.6$ |
| PLA ED | $8.0 \pm 1.0$ | $79.6 \pm 9.8$ | $2.6 \pm 0.2$ |
| PLA SD | $5.3 \pm 0.2$ | $44.1 \pm 1.8$ | $6.3 \pm 0.1$ |

In PLGA and PLGA-PEG nanoparticles, emulsion diffusion and solvent displacement resulted in nanoparticles with a similar drug load. In PLA nanoparticles, there was a substantial difference between the techniques (solvent displacement: 6.3 ± 0.1 µg/ doxorubicin/ mg nanoparticle, emulsion diffusion: 2.6 ± 0.2 µg doxorubicin/mg nanoparticle) (Table 2).

All nanoparticles displayed a similar drug release behaviour characterised by an initial burst release (Figure 3), which is in accordance to previous studies and caused by the release of drug adsorbed to the nanoparticles [Corrigan & Li, 2009]. PEGylated polymers may result in a more porous particle structure, which is caused by aqueous channels created by PEG chains and anticipated to further increase the initial burst release [Ruan & Feng, 2003]. However, we did not observe a particularly pronounced burst release in PLGA-PEG nanoparticles. A slight drop in the doxorubicin concentration was noticeable in the medium of the PLGA nanoparticles.

**Figure 3:**
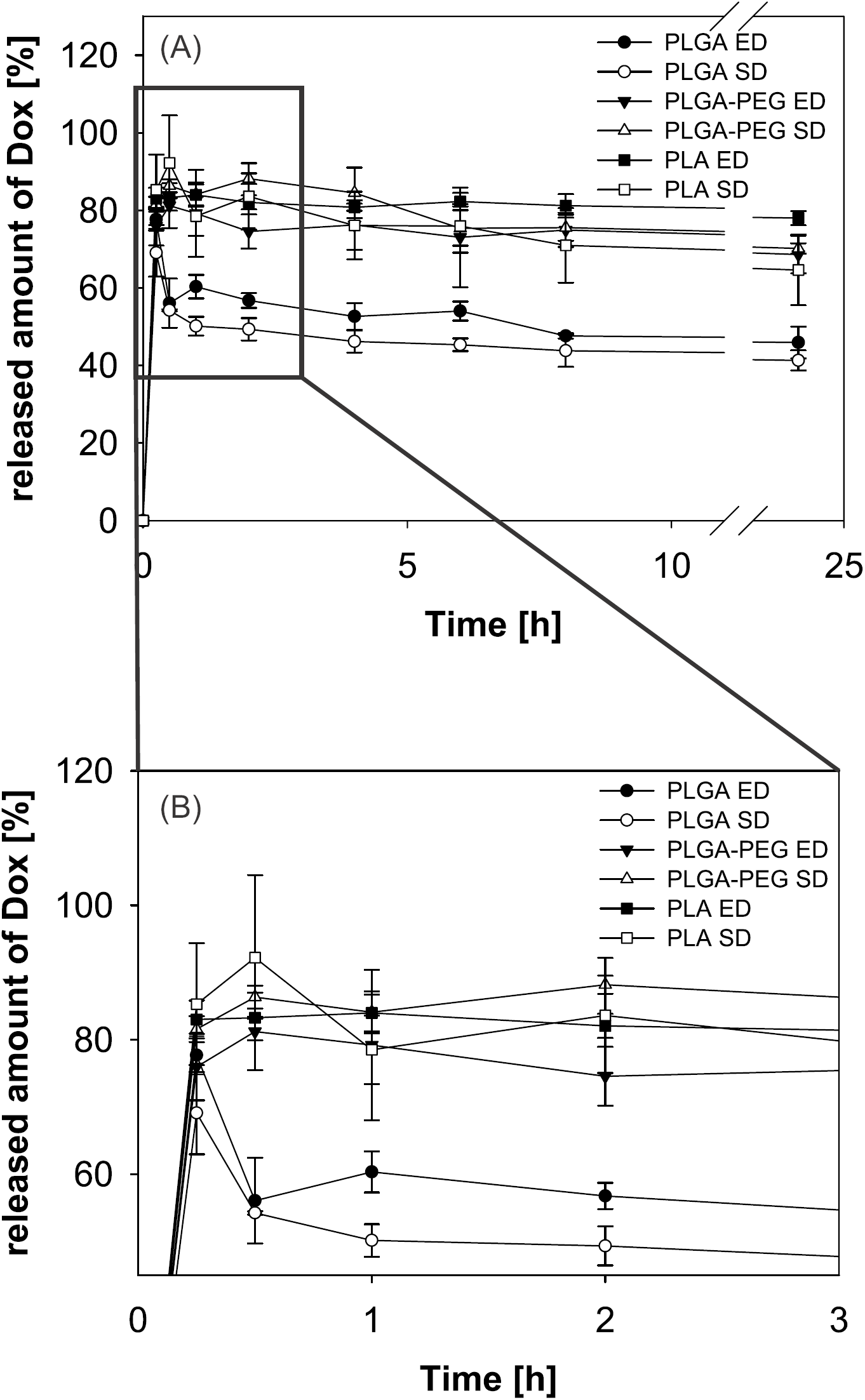
(A) Doxorubicin (Dox) release profiles for all nanoparticle systems using emulsion diffusion (ED) or solvent displacement (SD) preparation technique over 24 h. (B) Detailed section of the timeframe 0–3 h (data expressed as means ± SD, n = 3).

**Figure 4:**
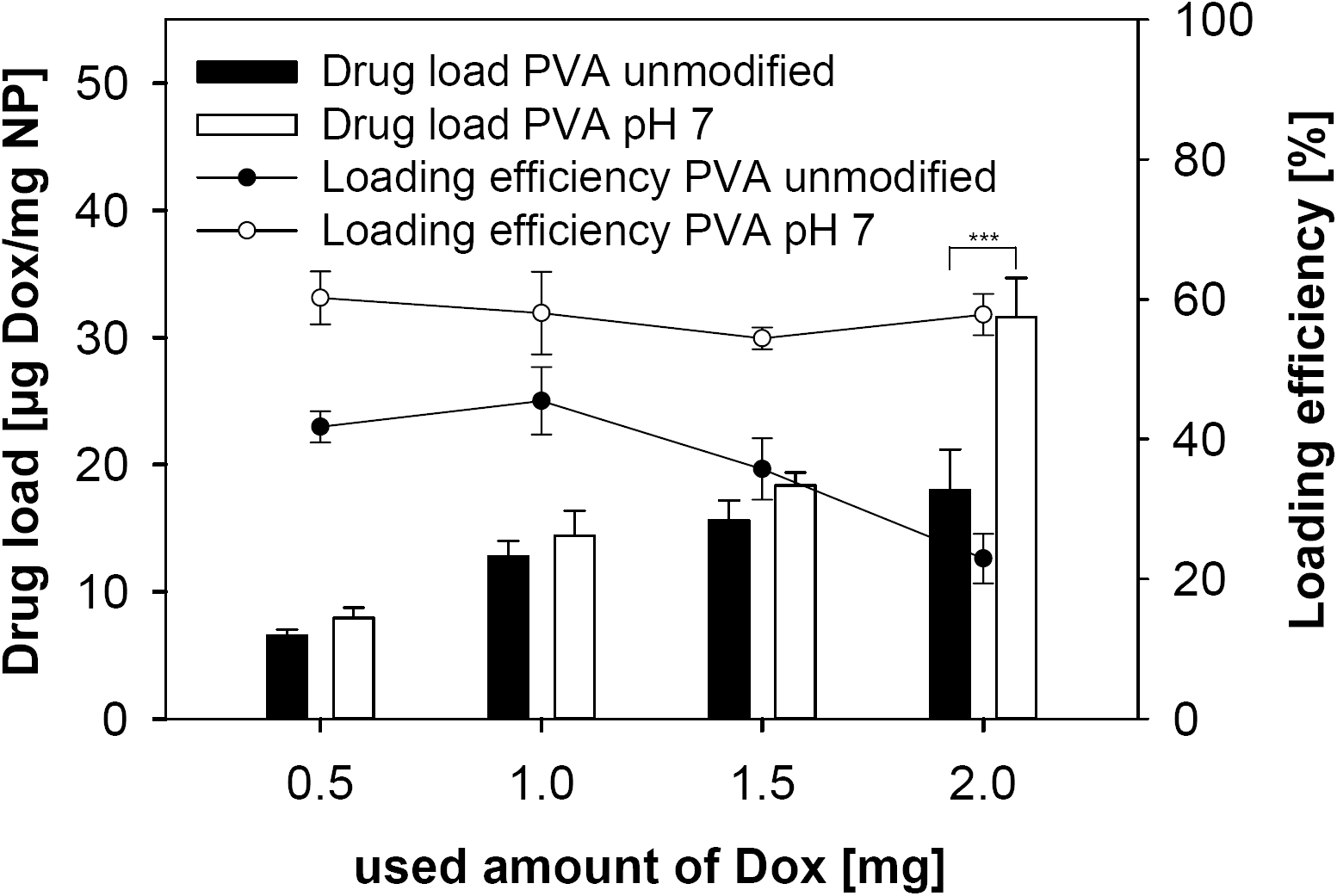
Doxorubicin (Dox) load and loading efficiency for PLGA nanoparticles (NPs) prepared using an unmodified PVA solution and a PVA solution adjusted to pH 7 (data expressed as means ± SD, n = 3).

This may be caused by a combination of doxorubicin adsorption to BSA, which was added to simulate the presence of plasma proteins, and slower doxorubicin release compared to the other nanoparticle systems.

Burst release suggests that a drug is primarily bound to the nanoparticle surface and not incorporated into the particle matrix. Such a burst release should be avoided, because it may result in drug release shortly after i.v. application before the nanoparticles reach the desired site of drug action (e.g. the tumour tissue) [Danhier et al., 2012].

To optimise the loading efficiency and drug release kinetics of PLGA nanoparticles the pH of the stabiliser solution used during nanoparticle preparation was increased to 7. At this pH value, doxorubicin exists in the more lipophilic deprotonated form. The use of PVA solution at pH 7 had no influence on the nanoparticle characteristics like particle diameter, PDI, and zeta potential (Table 3). However, loading efficiency and drug load increased. The drug load raised from 6.7 ± 0.3 µg doxorubicin/mg nanoparticle (44.8 ± 5.8% loading efficiency) without pH adjustment to 7.9 ± 0.8 µg doxorubicin/mg nanoparticle (60.2 ± 3.8% loading efficiency) at pH 7. By increasing the amount of doxorubicin to 2 mg, the drug load of PLGA nanoparticles could be further enhanced (non-adjusted pH: 18.0 ± 3.2 µg doxorubicin/mg nanoparticle; pH7: 31.6 ± 3.1 µg doxorubicin/mg nanoparticle, respectively) (Figure 4). Different amounts of doxorubicin did not change the loading efficiency at pH 7. Using aqueous solutions instead of methanol, we increased the doxorubicin amount to 5 mg and 7.5 mg doxorubicin per 50 mg PLGA. While 5 mg resulted in an increase of the drug load to of 52.5 ± 0.4 µg doxorubicin/mg nanoparticle, 7.5 mg doxorubicin did not result in a significant further increase (54.4 ± 3.4 µg doxorubicin/mg nanoparticle) (Figure 5A). A further increase of doxorubicin resulted in unstable nanoparticle systems, as indicated by increasing particle diameter and polydispersity index (Figure 5B). The loading efficiency for PLGA nanoparticles prepared at pH 7 with 5 mg doxorubicin was higher than this for nanoparticles manufactured with 7.5 mg doxorubicin (50.6 ± 0.6% and 33.9 ± 0.5%, respectively). In addition, PLGA nanoparticles prepared at pH 7 displayed a much more controlled and sustained doxorubicin release than PLGA nanoparticles prepared without pH adjustment (Figure 6). Hence, PLGA nanoparticles prepared at pH 7 with 5 mg doxorubicin were selected for cell culture experiments.

**Table 3:** Resulting particle diameter, PDI, and zeta potential (ZP) for PLGA nanoparticles prepared by an unmodified PVA solution and a PVA solution adjusted to pH 7 (data expressed as means ± SD, n = 3).

| PVA<br>solution | Diameter [nm] | PDI | ZP [mV] |
| --- | --- | --- | --- |
| unmod | $177.9 \pm 1.0$ | $0.039 \pm 0.031$ | $-41.6 \pm 2.0$ |
| pH 7 | $174.1 \pm 2.8$ | $0.057 \pm 0.030$ | $-43.8 \pm 3.7$ |

**Figure 5.**
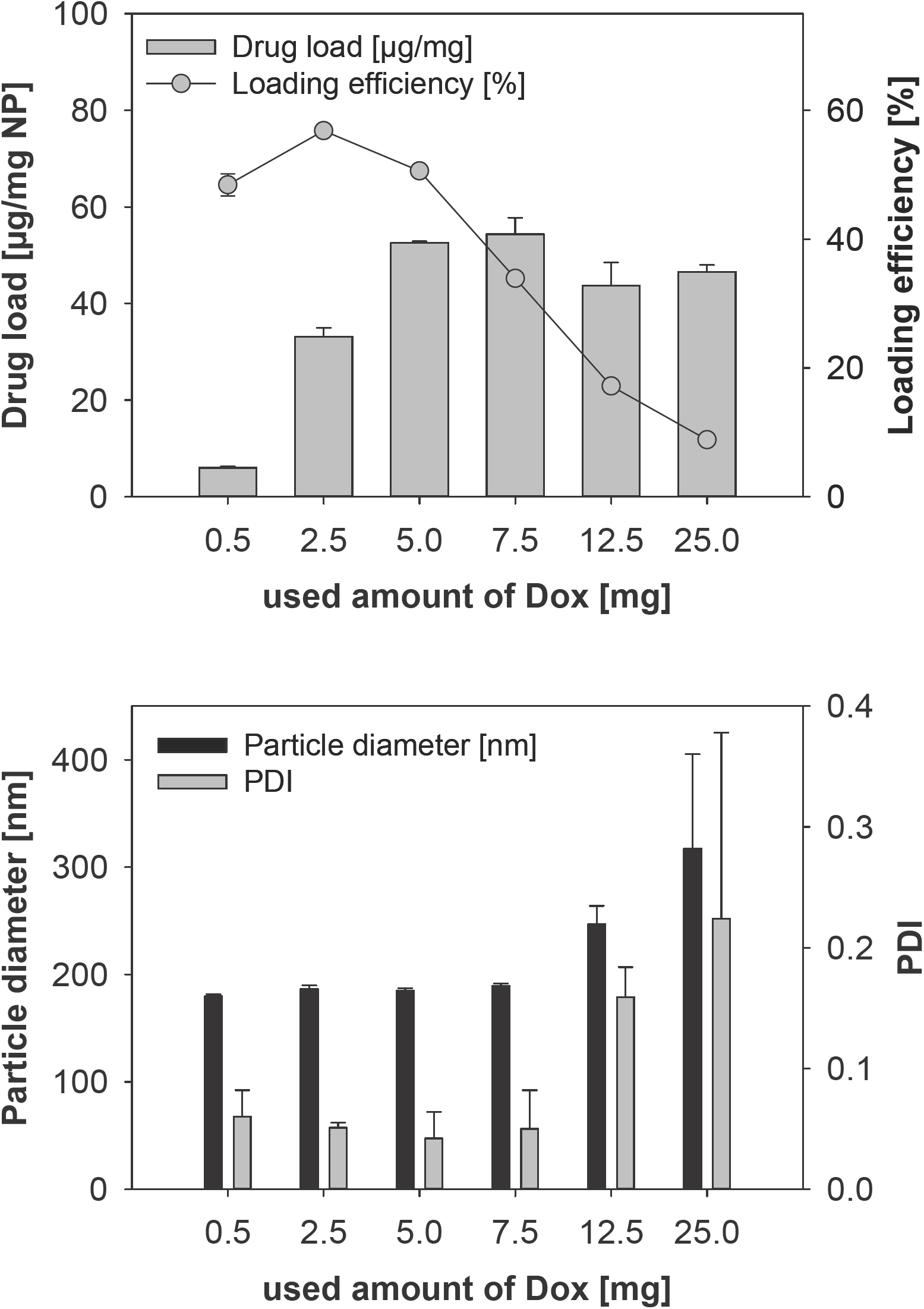
(A) Drug load and loading efficiencies as well as (B) particle diameter and PDI for different amounts of doxorubicin (Dox) used for the preparation of PLGA nanoparticles by emulsion diffusion technique (data expressed as means ± SD, n = 3).

**Figure 6.**
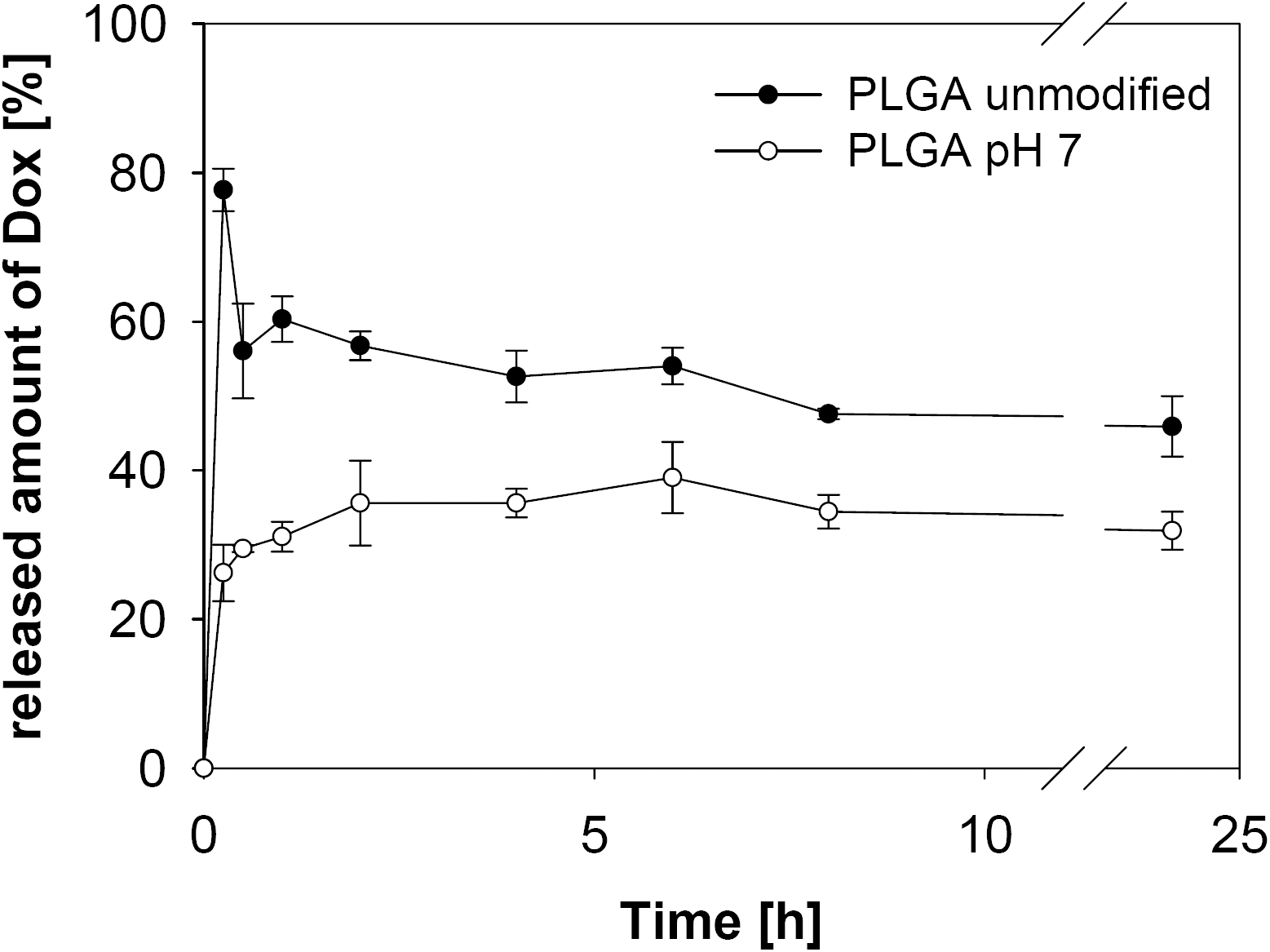
Release profiles of doxorubicin from PLGA nanoparticles prepared using an unmodified PVA solution and a PVA solution adjusted to pH 7 (data expressed as means ± SD, n = 3).

The different release kinetics from PLGA nanoparticles prepared at pH7, may be attributed to the higher lipophilicity of doxorubicin at this pH and in turn a stronger incorporation into the lipophilic PLGA nanoparticle matrix. This explanation is consistent with data showing that PLGA nanoparticle degradation is unlikely to occur in a 24 h timeframe [Li, 1999; Danhier et al., 2012].

### Nanoparticle efficacy in cell culture

Finally, the effects of doxorubicin-loaded PLA nanoparticles prepared by solvent displacement (because they were smaller and the drug load was higher compared to those prepared by emulsion diffusion), PLGA nanoparticles prepared by solvent displacement at uncontrolled pH and at pH7, and PLGA-PEG nanoparticles prepared by emulsion diffusion and solvent displacement were tested on the viability of the neuroblastoma cell line UKF-NB-3, its doxorubicin-adapted sub-line UKF-NB-3^r^DOX^20^, and its vincristine-resistant sub-line UKF-NB-3^r^VCR^1^.

In all three cell lines, PLA nanoparticles, PLGA nanoparticles prepared by solvent displacement at uncontrolled pH, and PLGA-PEG nanoparticles prepared by emulsion diffusion displayed reduced efficacy compared to doxorubicin solution (Figure 7). In contrast, PLGA nanoparticles prepared by solvent displacement at pH7 and PLGA-PEG nanoparticles prepared by solvent displacement were similarly active as free doxorubicin (Figure 7). The corresponding empty nanoparticles did not affect cell viability in the tested concentrations.

**Figure 7.**
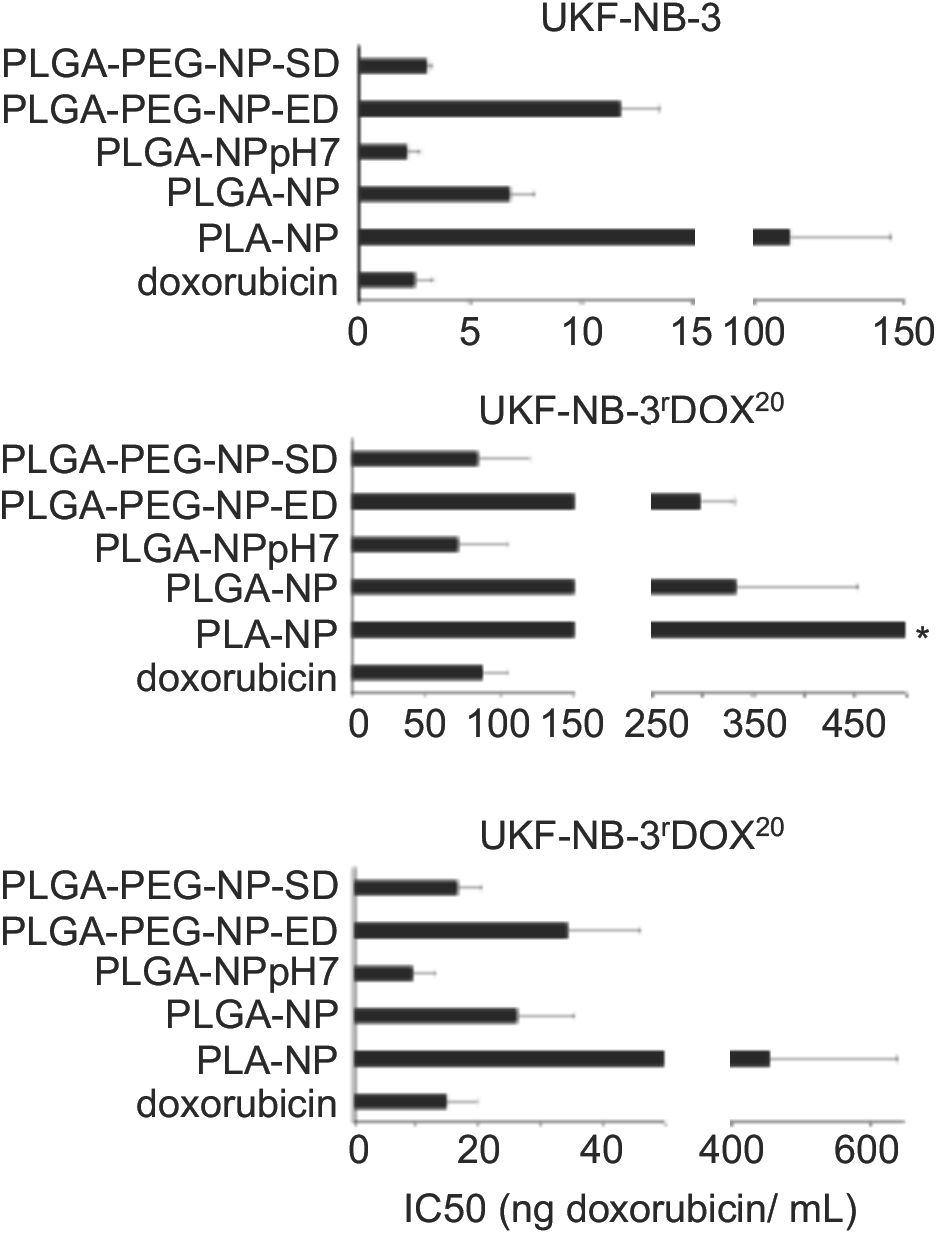
Doxorubicin concentrations that reduce neuroblastoma cell viability by 50% (IC50) when administered encapsulated into different nanoparticle preparations (PLA-NP, PLA nanoparticles prepared by solvent displacement; PLGA-NP, PLGA nanoparticles prepared by solvent displacement at non-adjusted pH; PLGA-NPpH7, PLGA nanoparticles prepared by solvent displacement at pH7; PLGA-PEG-ED, PLGA-PEG nanoparticles prepared by emulsion diffusion; PLGA-PEG-SD, PLGA-PEG nanoparticles prepared by solvent displacement) compared to doxorubicin solution (doxorubicin). Unloaded nanoparticles did not affect cell viability in the tested concentration range. * IC50 > 500 ng/mL

The main difference between the doxorubicin-loaded PLGA-PEG nanoparticles prepared by solvent displacement and the other preparations is the size. It is the only preparation in which nanoparticles have a size clearly smaller than 100 nm (72.6 ± 3.3 nm, Figure 1). This indicates that the cellular uptake of smaller nanoparticles is higher than that of larger nanoparticles, which is coherent with previous findings showing that cellular uptake of nanoparticles decreases with an increase of size [Salatin et al., 2015]. PLGA nanoparticles prepared by solvent displacement at pH7 displayed the highest drug load. Hence, their superior effects may be explained by an increased drug transport per nanoparticle into cancer cells.

Nano-sized drug carriers have been shown to bypass efflux-mediated drug resistance [Bar-Zeev et al., 2017]. This included various nanoparticle and liposome formulations of the ABCB1 substrate doxorubicin that were shown to modify the cellular uptake and intracellular distribution of doxorubicin resulting in enhanced effects against ABCB1-expressing cancer cells, when compared to free doxorubicin in solution [Thierry et al., 1993; Bennis et al., 1994; Wong et al., 2006; Prados et al., 2012; Oliveira et al., 2016; Maiti et al., 2018]. The doxorubicin-adapted UKF-NB-3 sub-line UKF-NB-3^r^DOX^20^ is characterised by high ABCB1 expression [Kotchetkov et al., 2005]. In addition, the vincristine-resistant resistant UKF-NB-3 sub-line UKF-NB-3^r^VCR^1^ displays cross-resistance to doxorubicin and becomes sensitised to doxorubicin by the specific ABCB1 inhibitor zosuquidar (Suppl. Figure 1). This indicates that drug resistance is at least in part mediated by ABCB1 in this cell line. However, free doxorubicin solution and doxorubicin bound to PLGA-PEG nanoparticles prepared by solvent displacement or PLGA nanoparticles prepared by solvent displacement at pH7 displayed similar efficacy in UKF-NB-3^r^DOX^20^ and UKF-NB-3^r^VCR^1^ cells (Figure 7). Hence, these drug carrier systems are not able to overcome transporter-mediated drug resistance.

## CONCLUSION

In this study, we synthesised a range of doxorubicin-loaded PLA- and PLGA-based nanoparticle systems using emulsion diffusion and solvent displacement approaches. Our results show that particle size, loading efficiency, and drug release kinetics can be controlled by the production procedure. Testing of the nanoparticle preparations in the neuroblastoma cell line UKF-NB-3 and its sub-lines with acquired resistance to doxorubicin or vincristine indicated that smaller nanoparticles and a high drug load result in nanoparticle preparations that have a similar efficacy at the cellular level as doxorubicin solution. Since nanoparticle preparations are known to have the capacity to improve the in vivo activity of anti-cancer drugs by tumour targeting through the EPR effect, this is an important step in the development of improved nanoparticle preparations

---

**Figure S1.**
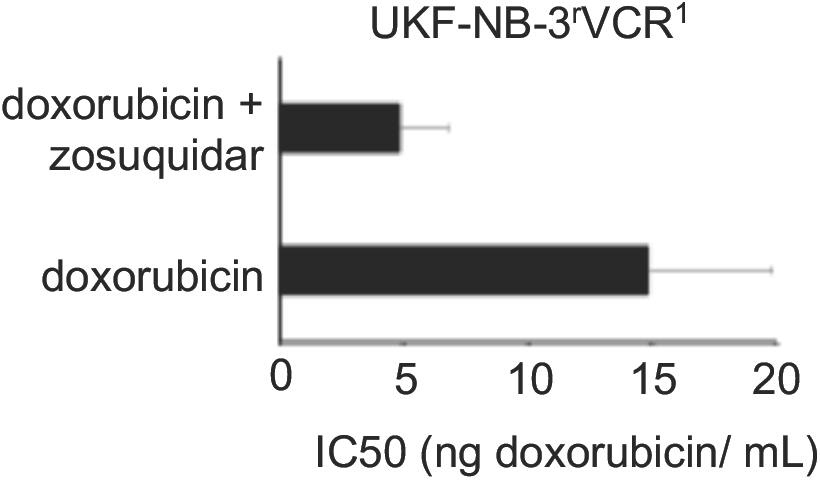
Doxorubicin concentrations that reduce UKF-NB-3^r^VCR^1^ viability by 50%455(IC50) in the absence or presence of the ABCB1 inhibitor zosuquidar (1µM). Zosuquidar did not affect cell viability when administered alone.

